# Slow and fast gamma oscillations show phase-amplitude coupling with distinct high-frequency bands in macaque primary visual cortex

**DOI:** 10.1101/2024.11.20.624422

**Authors:** Pooja Prabhu, Supratim Ray

**Affiliations:** Centre for Neuroscience, Indian Institute of Science, Bengaluru, India, 560012, Facsimile

**Keywords:** Gamma, Matching Pursuit, Phase-Amplitude Coupling, sharp transients

## Abstract

Gamma oscillations (25-70 Hz) can be induced in the visual cortex by presenting stimuli like gratings. Large stimuli produce two distinct gamma oscillations in primate primary visual cortex (V1) – slow (25-40 Hz) and fast (40-70 Hz), possibly due to different interneuronal networks. While fast-gamma has been shown to strongly lock spikes, slow-gamma does not, and hence its role in cortical processing is unclear. Here, we studied potential influence of gamma rhythms on neural activity using phase-amplitude coupling (PAC). We recorded spikes, local field potential and electrocorticogram (ECoG) from V1 of two adult female monkeys while presenting stimuli that simultaneously induced both gamma rhythms, and developed a novel method that reduces the influence of spike-related-transient on PAC. Interestingly, while fast-gamma showed coupling with frequencies above 150 Hz, reflecting spike-locking, slow-gamma showed PAC in a distinct frequency range between 80-150 Hz, which was especially prominent in ECoG. PAC varied with orientation and spatial frequency in the same way as power in the respective frequency bands, confirming dissociation in the coupling of the two gamma rhythms. Thus, fast-gamma could be more involved in spike-locking, while slow-gamma could represent a modulatory signal acting by amplitude modulation between 80-150 Hz at a more global scale.

**Significance Statement:** Gamma oscillations in the visual cortex can be induced by stimuli like gratings, producing two distinct gamma oscillations: slow (20-40 Hz) and fast (40-70 Hz). While fast-gamma strongly locks spikes, the role of slow gamma is unclear. Oscillations have been proposed to influence processing through phase-amplitude coupling (PAC). We recorded spikes, local field potential (LFP) and electrocorticogram (ECoG) from female monkeys and developed a new method to study PAC. While fast-gamma showed PAC with 150-500 Hz, reflecting spike-locking, we found PAC between slow-gamma and 80-150 Hz, which was especially strong in ECoG. The two PAC signatures varied differently with stimulus, reflecting distinct origins. Thus, while fast-gamma could lock spikes, slow-gamma could modulate amplitudes between 80-150 Hz at a global scale.

## Introduction

Gamma rhythm (25–70 Hz) observed in brain signals such as electroencephalogram (EEG) and Local Field Potential (LFP) has been linked to higher order cognitive processes such as perception, memory, and attention (1–3). Gamma has been thoroughly investigated in the primary visual cortex (V1) using visual stimuli such as bars and gratings (4, 5), and has been shown to depend on the attributes of gratings like size, orientation and spatial frequency (6, 7). In particular, large (full-screen) gratings produce two distinct gamma oscillations in V1 – slow (25-40 Hz) and fast (40-70 Hz), which are tuned to different orientations and have distinct temporal evolution profiles (7). Slow-gamma overlaps with the beta (16-30 Hz) range, but we term it differently to differentiate it from the classical beta rhythm which is prominent in sensory-motor areas and is modulated by motor planning (8). In particular, while field-field coherence shows stronger coherence and a slower decay with increased inter-electrode distance for slow gamma compared to fast gamma, spike-field coherence shows stronger coupling of spikes with fast-gamma (7). Therefore, how slow-gamma affects neural activity remains unclear.

Studies on rodents suggest that distinct inter-neuronal networks could produce distinct gamma waves (9, 10). GABAergic interneurons show various biochemical, morphological, and electrophysiological characteristics in numerous brain circuits (11), which could be involved in the generation of oscillatory activity (12). For example, somatostatin-expressing (SOM) interneurons are crucial in producing oscillations in the slow gamma (20–40 Hz) range in the neocortex (9, 10), while parvalbumin-expressing (PV) interneurons are crucial in producing both theta (4–8 Hz) (13, 14) and fast gamma (30–80 Hz) rhythms (9, 15). Interestingly, these interneurons target different parts of the neuron, with SOM interneurons preferentially targeting dendrites while PV interneurons targeting the soma, leading to differences in the way they affect neural responses (16–18).

One way an oscillation can influence neural activity is through cross-frequency-coupling, where the phase/amplitude of one oscillation is coupled to the amplitude of another (for example, phase-amplitude coupling (PAC; (19, 20)). Previous studies on macaques have investigated PAC between slow waves and gamma during sleep in the hippocampus (21), in directional communication in the auditory cortex (22), and between beta and gamma in directional communication in V1 (23). However, in most of these studies, gamma oscillations were endogenously produced, not stimulus-induced. Further, since the two gamma rhythms may operate on different spatial scales, it is important to study PAC across scales.

Here, we examined PAC in simultaneously recorded LFP and ECoG from V1 of macaques while they viewed large gratings optimized to simultaneously produce strong slow and fast gamma oscillations. Since spikes are associated with an increase in power in the high-gamma range (> 80 Hz; (24, 25)), and fast-gamma strongly lock spikes, we expected strong PAC between fast-gamma and high-gamma. However, the spike-related-transient in the LFP is well represented as a negative Gaussian (26, 27), and such non-sinusoidal oscillations or sharp transients produce unintuitive phase and amplitude estimates (28, 29) and cause false increases in the amplitude of high-frequency oscillations (28), leading to spurious PAC. We therefore used two different approaches to reduce this component and then studied PAC between oscillations of phase up to 80 Hz (which covers both gamma bands) and amplitudes up to 500 Hz.

## Results

We simultaneously recorded spikes, LFP and ECoG using a customized hybrid electrode array from V1 of two monkeys while they viewed full screen gratings of varying orientations and spatial frequencies (see (30) and Methods for more details). These stimuli produced two distinct gamma waves with some variation in amplitude and centre frequency with orientation and spatial frequency in both LFP (Supplementary Figure 1) and ECoG (not shown; ECoG and LFP tuning was very similar; see (30)). The power spectral density (PSD) averaged across all stimulus conditions showed two peaks between 25-40 Hz and 40-70 Hz, which were chosen as the frequency bands for slow and fast gamma for further analysis (Figure 1).

**Figure 1:**
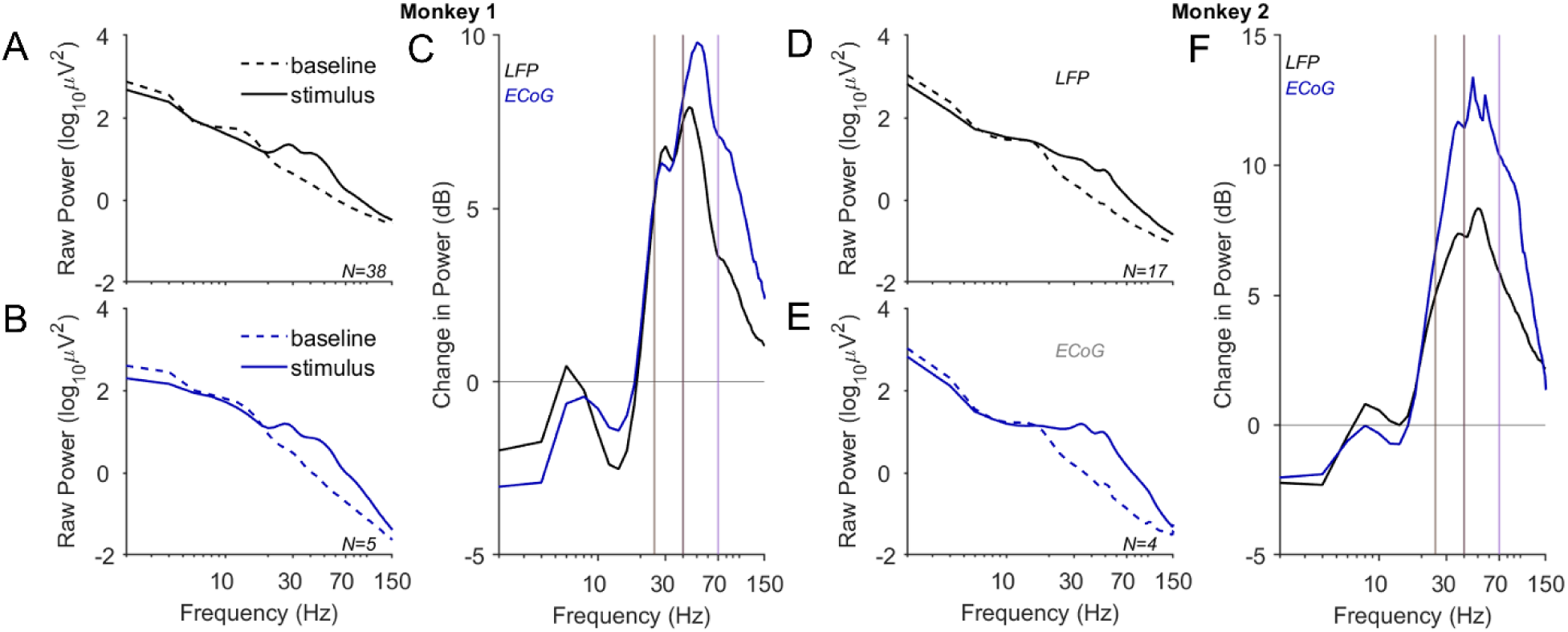
Raw and change in power across all the conditions in LFP and ECoG. The raw power for baseline (−500 ms to 0) and stimulus (250 ms to 750 ms) period is calculated across all the stimulus conditions in LFP (A and D) and ECoG (B and E) for Monkey 1 and 2. The change in power for Monkey 1 and Monkey 2 shows the peak in slow gamma (25– 40 Hz; brown vertical lines) and fast gamma (40-70 Hz; violet vertical lines) range for LFP and ECoG for both monkeys (C and F).

### Spike and LFP are predominantly coupled in a fast gamma range

To test how these oscillations influence spiking activity, we computed the spike-triggered-average (STA), where we took segments of LFP centered around spikes and averaged these segments (Figure 2A-B; magenta traces). These traces show an oscillatory pattern with a period of ∼50 ms with a trough near spike onset (t=0) and another transient structure of a shorter duration (a few ms), which we call the “spike-related-transient”, which could be due to the synaptic activity that produces the spike as well as the low-frequency component of the action potential itself (“spike-bleed-through”), as discussed in detail elsewhere (26, 27, 31)). The spike-related-transient cannot be accurately represented using classical spectral methods that decompose the signal into long sinusoids, but can be decomposed using the Matching Pursuit (MP) algorithm which has delta functions as well as sharp Gaussians as potential basis function to represent the transient activity (see Figure 2 of (31) and Figure 4 of (27) for MP based decomposition of the STA). In particular, there is a sharp negative deflection around spike onset which is captured by a negative Gaussian. Such a structure is likely to produce erroneous PAC because the Fourier decomposition of such a Gaussian yields a large set of sinusoids whose troughs are aligned, and hence produces artifactual coupling across multiple sinusoids. We therefore decomposed the LFP signal using MP and removed all basis functions (called atoms) whose centre frequency was zero (which include delta functions and Gaussians). STA after MP based removal showed a slight modification in the spike-related-transient (Figure 2A-B; green trace), but not a complete removal because part of this spike-related-transient was represented by high-frequency sinusoids, not Gaussians. Another approach to remove the effect of the spike-related-transient is to use LFP from a neighbouring electrode (32). This approach reduced the transient, allowing a better representation of the slow oscillatory component (Figure 2A-B; yellow traces).

**Figure 2:**
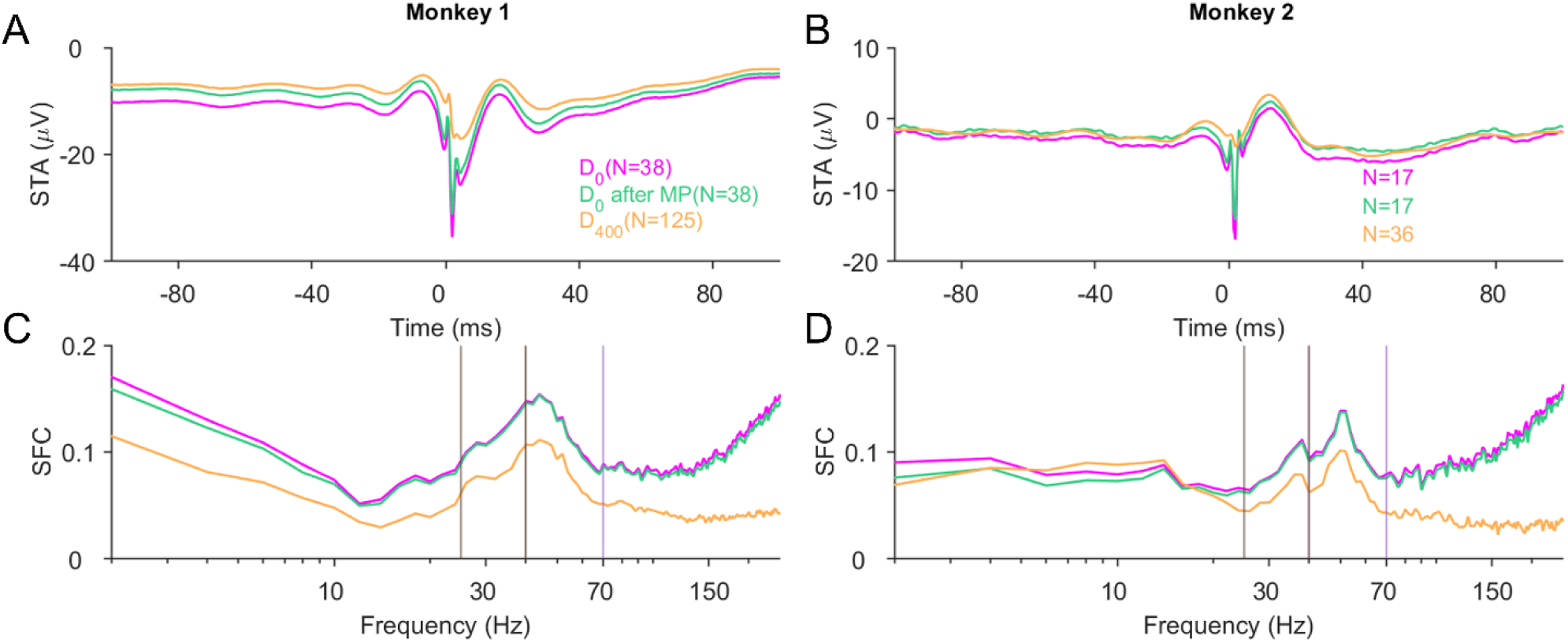
Spike Triggered Average (STA) and Spike-LFP Coherence (SFC) across all stimulus conditions in LFP. The averaged STA (A-B) curve (pink) shows the negative deflection (or negative Gaussian). After MP based removal of non-sinusoidal components (brown curve), there was a slight reduction in negative deflection, and by considering spikes and LFP from neighboring electrodes (mustard curve), the negative Gaussian component was further reduced. The SFC (C-D) shows a strong peak in the fast gamma range but a smaller peak in the slow gamma range, suggesting preferential coupling of spikes with fast gamma. When spikes and LFP are recorded from the same electrode (pink and magenta traces), there is high SFC at frequencies above 100 Hz because spikes produce spike-related-transients in the LFP with energy at these frequencies. Removal of the non-sinusoidal component only leads to a slight reduction in SFC (magenta versus green traces). Computing SFC with spikes and LFPs from neighboring electrodes leads to a substantial reduction of high-frequency SFC, although the coupling to the gamma rhythm remains prominent.

To find the frequencies of these oscillations, we computed the spike-field-coherence (SFC), which can be shown as the Fourier transform of the STA with appropriate normalization across frequencies. The SFC showed a more prominent peak at fast-gamma compared to slow-gamma, indicating preferential coupling of spikes with fast-gamma. The spike-related-transient, which has high-frequency components, led to an artifactually high value of SFC at frequencies above ∼150 Hz (Figure 2C-D; magenta traces), which was considerably reduced when LFP was taken from a neighbouring electrode (yellow traces). The effect of the Gaussian removal led to a minor reduction of SFC, mainly at frequencies below 10 Hz (compare magenta versus green traces in Figure 2C-D).

### PAC reveals distinct coupling of slow and fast gamma

We computed PAC by filtering signals in different frequency bands and computing their instantaneous phase and amplitudes using Hilbert transform (see Methods and (19)). As before, we either used the raw LFP (first column in Figure 3A for Monkey 1 and 3B for Monkey 2) or the LFP after removal of Gaussian components (middle columns). We also tested PAC after computing phases and amplitudes from neighbouring electrodes (last column).

**Figure 3:**
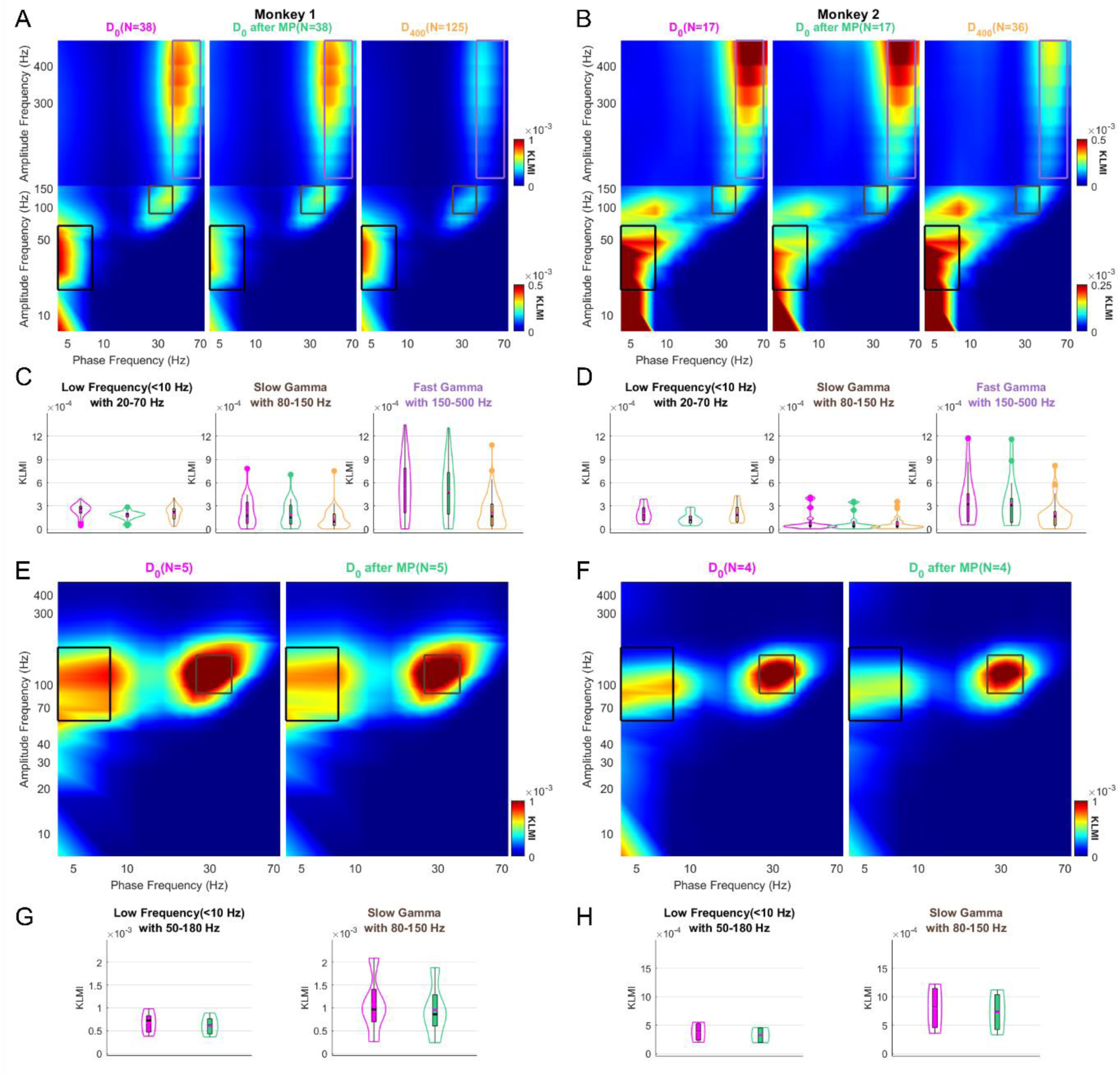
Effect of Matching Pursuit and neighboring electrodes on PAC in LFP and ECoG data. A-B) PAC calculated for three conditions: using raw LFP (D_0_; left column), after removing non-sinusoidal components using MP (D_0_ after MP; middle column), and when phase and amplitude are calculated for neighboring electrodes separated by 400 microns (D_400_; right column). PAC predominantly showed three clusters in low frequency (black box), slow gamma (brown box), and fast gamma (violet box) range for LFP data. C-D) Average KLMI values computed by averaging the values in the three clusters shown in A-B. D-H) Same as A-D but for ECoG data. The neighboring condition is not used since the ECoG electrodes were separated by 10 mm.

We found strong PAC between phase of low-frequency (4-8 Hz) and amplitude between 20-70 Hz (indicated with a black box in Figure 3A and B). This coupling extended to lower amplitude frequencies as well, especially in Monkey 2, but we did not consider this range for analysis because PAC can be spurious for amplitude frequencies lower than phase frequencies (33). Even though the Gaussian removal process had a minor effect on STA and SFC, it resulted in a substantial reduction in PAC for this low-frequency cluster (compare the boxes in the first and middle panels). Average PAC values (or KLMI) within this low-frequency (Figure 3; C and D, first column) reduced by 21% and 30% for the two monkeys (t=3.64, p=3.9×10^-3^ for Monkey 1 and t=3.61, p=4×10^-3^ for Monkey 2). This suggests that even small transient non-sinusoidal structures can impact PAC analysis. However, this low-frequency PAC did not change significantly (t=0.47, p=0.63 in Monkey 1 and t=0.058, p=0.954 in Monkey 2) when the phase and amplitudes were computed from neighbouring electrodes (compare first versus last columns in Figure 3A and 3B).

On the other hand, fast-gamma predominantly showed PAC with a broad frequency range between 150-500 Hz (violet box). This is consistent with the previous observation that spikes get locked preferentially to the fast-gamma and the spike-related-transient has prominent energy above 150 Hz (Figure 2C and 2D). Consistent with the SFC results, removal of the negative Gaussian had a small effect (7.51% and 7.44% reduction for the two monkeys), although the difference was still highly significant using a paired t-test since there was always a reduction in PAC (t=44.23, p=1.75×10^-132^ for Monkey 1 and t=30.48, p=2.56×10^-93^ for Monkey 2). On the other hand, choosing different electrodes for phase and amplitude lead to a substantial reduction in PAC (compare middle and last columns). Choosing neighboring electrodes showed the significantly more reduction in PAC in both the monkeys (60% reduction, t= 24.21, p=1.53×10^-90^ in Monkey 1 and 47%, t=19.53, p=5.93×10^-66^ in Monkey 2; note that t-statistics and p-values are not directly comparable with the previous comparison because here we performed an unpaired t-test).

Interestingly, we found that slow-gamma showed PAC with frequencies between 80-150 Hz (brown patch), even though such PAC was not observed with fast-gamma. Similar to fast-gamma PAC, removing the Gaussian component had a modest effect (reduction of 12% and 13%; t=23.87 and p=6.94×10^-25^ for Monkey 1 and t=19.41 and p=1.18×10^-21^ for Monkey 2), but computing phase and amplitude from neighbouring electrodes showed substantial reduction (42% reduction, t=8.58, p=6.96×10^-13^ in Monkey 1 and 22.15%, t=4.01, p=1.34×10^-4^ in Monkey 2).

While the slow-gamma and 80-150 Hz PAC was weaker than fast-gamma and 150-500 Hz PAC (note difference in scale in the upper and lower PAC plots) in LFP data, we observed a different trend for ECoG electrodes (Figure 3; E and F; neighboring electrode condition was not considered). For ECoG electrodes, slow-gamma and 80-150 Hz PAC was the strongest, followed by low-frequency and 50-180 Hz, while the fast-gamma with 150-500 Hz PAC was negligible, even though fast-gamma was observed in the PSDs (Figure 1). Potential implications of this difference in PAC between LFP and ECoG signals is discussed later.

The phase-mean amplitude distributions showed that for both PACs, the amplitudes peaked near the trough of the corresponding rhythm (defined as 180 degrees; Supplementary Figure 2). For Monkey 1, for electrodes that showed significant PAC, amplitudes between 80-150 Hz peaked at mean phase of 165 ° (N=32) of slow gamma, while amplitudes between 150-500 Hz peaked at mean phase of 154° (N=22) of fast gamma. For Monkey 2, the corresponding values were 232° (N=10) and 195°(N=9).

### Orientation tuning for PAC follows the trends in power

In the LFP data, the 80-150 Hz band that coupled with slow-gamma appeared at the tail end of the broader 150-500 Hz band which showed PAC with fast-gamma. To test whether the coupling between 80-150 Hz was due to coupling with a different frequency band (slow-gamma) as compared to a residue of a broad-frequency coupling with fast-gamma, we computed PAC individually for stimuli of different orientations, since power of slow and fast gamma changed differently with orientation, with strongest slow gamma at 0° and 45° for the two monkeys and strongest fast gamma at 90° and 67.5° (Supplementary Figure 1). Interestingly, the PAC values (KLMI) within slow-gamma (brown curves in Figure 4; C and E) and fast-gamma range (violet curve in Figure 4; C and E) averaged across electrodes (38 and 17 electrodes for Monkey 1 and Monkey 2, respectively) changed with orientation in the same way as power changed with orientation in the two gamma bands (Figure 4; D and F). The Spearman correlation between PAC and power was high for both gamma bands: slow gamma PAC and slow gamma power (r=0.78, p=0.027 for Monkey 1 and r=0.71, p=0.05 for Monkey 2), fast gamma PAC and fast gamma power (r=0.90, p=0.004 for Monkey 1 and r=0.95, p=0.001 for Monkey 2), suggesting that PAC followed similar trend as that of change in power in LFP. In contrast, PAC and power from different frequency bands were not significant for either monkey. Similar trends were obtained for slow gamma in ECoG as well (Figure 4G-L).

**Figure 4:**
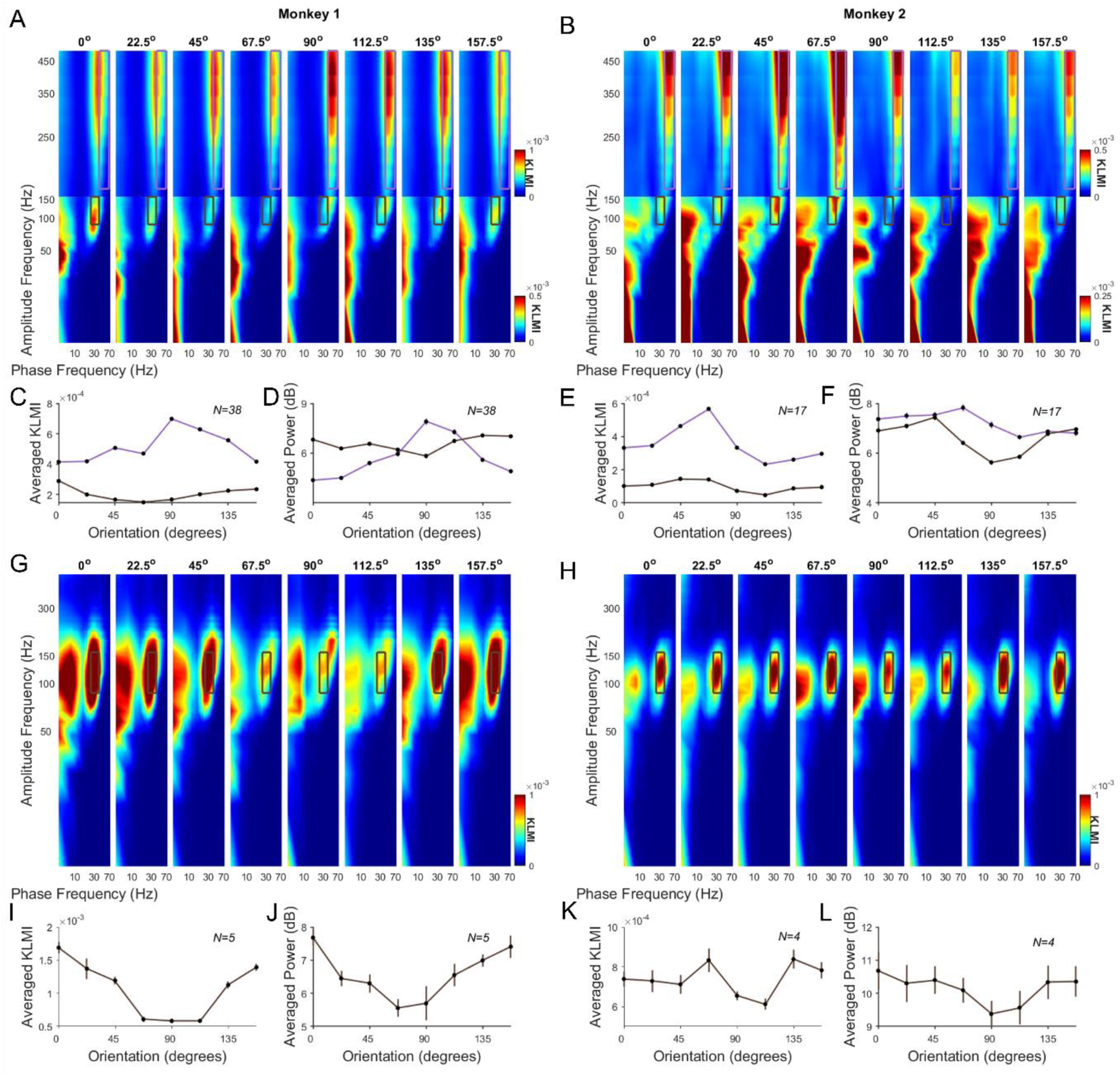
Orientation tuning of slow gamma and fast gamma PAC in LFP and ECoG after removal of sharp transients. (A-B) PAC plots showing slow and fast gamma coupling to different high frequency ranges across different orientations. The PAC values (or KLMI) within slow gamma (brown curve) and fast gamma range (violet curve) were averaged across electrodes (38 and 17 electrodes for Monkey 1 and Monkey 2, respectively) to get tuning curves (C and E). The change in power of LFP for each orientation was computed for different orientations in slow gamma and fast gamma range (D and F). G-L) Same for ECoG data, for slow gamma PAC.

## Discussion

We observed strong PAC between theta (4-8 Hz) and a broad frequency range including the gamma band (30-100 Hz), but it was strongly attenuated once the spike-related transient was accounted for. We also observed fast gamma (40-70 Hz) PAC with frequencies above 150-500 Hz, potentially reflecting spike-locking. In addition, we found that simultaneously induced slow gamma (25-40 Hz) coupled preferentially to a distinct frequency range between 80-150 Hz. This slow gamma PAC was even more prominent in simultaneously recorded ECoG, for which fast gamma did not show any PAC. Slow and fast gamma power varied differently with orientation and spatial frequency, and PAC followed the same trend as power, reflecting distinct origins of the two PAC signatures.

### Neural mechanisms underlying slow and fast gamma PAC

Rodent studies have shown that somatostatin-expressing (SOM) interneurons are crucial in producing oscillations in the slow gamma range in the neocortex (9), while parvalbumin-expressing (PV) interneurons were crucial in producing both theta (4–8 Hz) (13, 14) and fast gamma (30–80 Hz) rhythms (9, 15). These interneurons have different morphological characteristics, electrophysiological properties and synaptic targets.

The PAC results shown here are consistent with the action of these two interneuronal classes on pyramidal neurons. SOM interneurons preferentially target the dendrites of pyramidal cells, and show broader dendritic arborization, enabling them to pick up inputs from broader range of sources, which could make slow gamma a global modulatory signal. On the other hand, PV interneurons preferentially target the soma of pyramidal cells and pick up inputs from localized sources (compact dendritic arborization), which makes fast gamma a local modulatory signal. PV and SOM interneurons synapse either to perisomatic-targeting or dendritic targeting regions making them deep or more superficial, respectively. Dendritic synaptic or action potentials, integrated over a broader range, are likely to be broader than somatic potentials (17), which could explain more power at relatively lower frequencies (80-150 Hz) in case of slow-gamma compared to fast-gamma PAC. This is also consistent with the observation that slow-gamma PAC with 80-150 Hz is more prominent in ECoG, because dendritic synaptic/spiking is likely to be more superficial, ECoG could be picking up more surface level signal when compared to LFP.

### Spurious PAC

While PAC examines the link between the phase and amplitude of various frequency bands in the signal, SFC evaluates the consistency of spike timing relative to the phase of the signal at different frequencies. Non-sinusoidal artifacts like Gaussian component modify the waveform shape and may introduce harmonics (34). Altered waveform shape, harmonic contamination and the non-linear effects are some of reasons to spurious PAC estimates.

Spurious relationship between spiking activity and LFP can arise due to the presence of “spike-related-transient” in the LFP, which is due to synaptic activity that generates the action potential as well as low-frequency component of the action potential itself (i.e., spike “bleed-through”) (27). Such contamination is readily evident if the LFP and spikes are recorded from the same electrode, but also observed if the recording LFP electrode is within a few hundred microns from the spike electrode (35). One approach is therefore to consider LFP electrodes beyond 200μm of the spike electrode (32), although that does not completely eliminate potential artifacts (Ray, 2015). Several approaches are used in removing these transients like estimating the spike-related-transient through matching pursuit algorithm (31) or using Bayesian inference (36), designing an optimal linear filter to compute the influence of spike on LFP (37) and adaptive artifact removal by decomposing individual action potentials into their frequency components and then removing from LFP (38).

These spike-related transients have a sharp gaussian-like component, which is not well represented by a sinusoidal signal. Further, even rhythmic components in a signal are often non-sinusoidal in shape, such as mu rhythm (8-10 Hz; (39)), beta rhythm (13-20 Hz)(40) and gamma rhythm (41). The sharp deflection of non-sinusoidal component constitutes a low frequency component which couples with high frequency amplitude peak contained in sharp deflection (42). When such data are filtered to get PAC, the phase at the frequency of occurrence of the potentials align with the amplitude peak, introducing artefactual or spurious PAC.

In literature, spurious PAC has been dealt differently. Soulat and colleagues (43) used a state-space model to better estimate the component oscillations before performing PAC. Velarde and colleagues (44) developed a time-Locked Index to quantify the harmonics generated by non-sinusoidal components which produce strong artifactual PAC. Jurkiewicz and colleagues (45) used an extended version of modulation Index used earlier (20) to eliminate the spurious coupling. This approach used the time-frequency representation of signal instead of filtering the signal to certain band-pass frequencies. In addition, the spurious PAC can be caused by insufficient epoch length, improper band pass filtering settings and improper surrogate methods (46). In our study, we have tackled spurious PAC by eliminating non-sinusoidal component directly using MP and also by performing inter-electrode PAC separated by 400μm.

We observed strong attenuation of PAC between theta (4-8 Hz) and a broad frequency range including the gamma band (30-100 Hz) after the removal of Gaussian component. However, we note that theta-gamma coupling has been mainly studied during encoding and retrieval in hippocampus (47, 48), which has strong theta oscillations, unlike V1 where the low-frequency rhythms get suppressed due to visual stimulation. Therefore, the theta-gamma coupling reported here (which is prone to the spike-transient artifact) could be very different from the one in hippocampus and other non-visual areas.

While our approach reduced theta-gamma PAC substantially, it had only a minor effect on SFC. This is not surprising, since SFC and PAC are very different measures. SFC is an estimate of the consistency of spike-timing relative to the phase at a given frequency, while PAC estimates the coupling between phase of low frequency with amplitude of high frequency (19, 20). One of the reasons behind a small effect of MP based removal of spike-transient on SFC could be because the removed component itself is small, which does not have a big effect at any individual frequency, but can have a more substantial effect when two frequencies are simultaneously compared, as in PAC.

### Functional role of PAC

PAC is thought to play a role in the dynamic coordination of brain circuits and systems. Studies have shown that PAC is useful in top-down processing of information processing (49) and information integration (50, 51). PAC between gamma and high gamma may be facilitating integration of top-down expectations with bottom-up sensory input (52). Our results suggest a more modest interpretation that does not involve a functional role of PAC. First, theta-gamma PAC, at least in part, could be an artifact due to the spike-related-transient, since removing this component substantially reduced PAC. Second, our results could be explained by locking of spikes to the trough of the gamma rhythms, since for both gamma rhythms, the largest amplitude was near the trough of the rhythm. As explained previously, potential differences in the shape of the synaptic/spiking activity (dendritic versus somatic) that was associated with slow versus fast gamma could potentially explain differential coupling to 80-150 Hz and 150-500 Hz bands, even though this type of PAC may not have any functional role. Therefore, our results suggest more caution while interpreting PAC results. On the other hand, differences in PAC for different gamma rhythms and scales could provide important clues about the underlying circuitry of cortex.

## Materials and Methods

### Animal Recordings

Two adult female monkeys (*Macaca radiata*) weighing 3.3 and 4 kg were used in the experiments. The protocols used for the animals were approved by the Committee for the Purpose of Control and Supervision of Experiments on Animals (CPCSEA) and the Institutional Animal Ethics Committee (IAEC) of the Indian Institute of Science. The experiments involved the simultaneous recording of spikes, LFP, and ECoG signals from the monkeys. The left cerebral hemispheres of the monkey were implanted with a specially designed hybrid array after it had learned the fixation task. Blackrock Microsystems manufactured the connector that connected the 3 × 3 ECoG electrodes (Ad-tech Medical Instrument) to the 9 × 9 microelectrodes in this hybrid array. Platinum discs with an exposed diameter of 2.3 mm and an inter-electrode center-to-center distance of 10 mm served as the ECoG electrodes. The microelectrodes measured 1 mm in length and 400 μm in spacing. The ECoG sheet was introduced subdurally so that the previously created silastic gap between four ECoG electrodes was aligned with the durotomy after a major craniotomy and a smaller durotomy. The electrode array was implanted under general anesthesia. Finally, the microelectrode array was placed into the opening at a distance of around 10-15 mm from both the midline and the occipital ridge (for a detailed explanation, refer to (7, 30)).

### Experimental Setup and Behavioral Task

The monkeys’ heads were fixed by the headpost while they performed the behavioral activity, sitting on a chair inside a Faraday cage enclosure to shield them from outside electrical noise. The visual fixation task was carried out by monkeys, wherein they had to maintain eye contact with a small fixation spot (0.05° or 0.1°) displayed in the middle of an LCD monitor screen that had been gamma-corrected within 2° of the fixation spot. The display was positioned 50 centimeters away from the subjects’ eyes, with full-screen gratings covering approximately 56° and 33° of the visual field in width and height.

The behavioral task began with the monkey holding and maintaining fixation. Following a 1000 ms initial blank phase, two to three stimuli were displayed one after the other for 800 ms each, separated by 700 ms between each stimulus. If the monkeys’ fixation remained constant during the trial, they got a drop of juice as a reward. Fullscreen static gratings at full contrast and one of eight orientations (0°, 22.5°, 45°, 67.5°, 90°, 112.5°, 135°, and 157.5°) and five spatial frequencies (0.5, 1, 2, 4, and 8 cpd) were presented pseudo-randomly. For Monkey 1 and Monkey 2, respectively, the average number of repetitions for each orientation condition and preferred spatial frequency was 33 (range 28 to 36) and 42 (range 37 to 45).

### Electrode Selection

Similar to our earlier research (7, 30), we limited the electrodes used for additional analysis to those whose receptive Field (RF) estimates were steady over the course of many days (SD less than 0.1°). This resulted in 77 and 32 LFP electrodes as well as 5 and 4 ECoG electrodes from Monkey 1 and 2.

Using Spikesort (53) (http://www.smithlab.net/spikesort.html), spike sorting was carried out on the chosen LFP electrodes. We determined the trial averaged change in firing rate in the 250–750 ms period from baseline (−500 to 0 ms) across stimulus conditions, as well as the signal-to-noise ratio for the sorted units. We selected units with a firing rate above 2 and a signal-to-noise ratio above 1.5, yielding 38 (52 units) and 17 (18 units) spike electrodes from Monkey 1 and Monkey 2, respectively.

### Data Analysis

Custom MATLAB algorithms were used to examine all of the data. The Matlab scripts used in the study are available as a Toolbox (https://github.com/poojaprabhu9/CrossFrequencyCouplingProject.git). For Monkeys 1 and 2, the baseline period was set between −500 and 0 ms (0 denotes the start of the stimulus), and the stimulus period was selected between 250 and 750 ms. Spike Field Coherence (SFC), spike-triggered Average (STA), and Power Spectral Density (PSD) were computed using the multitaper method implemented in Chronux 2.0 (54), an open-source data analysis toolbox available at http://chronux.org.

#### Power Spectral Density (PSD)

PSD was calculated using the multitaper method using 3 tapers for the baseline period and stimulus period, yielding a frequency resolution of 2 Hz. The PSD for each electrode was determined by independently calculating power for each trial and then averaging them. Change in power for each stimulus condition was calculated as follows: Power_i_= 10(log_10_ ST_i_ - BL_ave_), where ST_i_ is the power summed across the frequency range of interest for each of the gamma rhythms for stimulus condition *i* and BL_ave_ is the baseline power averaged across conditions (BL_ave_= average(log_10_ BL*i*)) (7).

#### Spike Field Coherence (SFC)

Using the Chronux toolbox, SFC was computed using multitaper method using a single taper for spike-LFP coherence in the stimulus period. Coherency was calculated on the data obtained by pooling across all five spatial frequencies and eight orientations. SFC was also calculated on LFP data after matching pursuit based removal of non-sinusoidal components.

#### Spike Triggered Average (STA)

Using the Chronux toolbox, we calculated the STA to investigate the relationship between spikes and LFP. STA was computed by selecting the segment from the LFP signal centered on each spike (100 ms of either side of the spike) and averaging over all segments within the stimulus period.

#### Phase Amplitude Coupling (PAC)

PAC is a neural oscillation phenomenon where the amplitude of higher-frequency oscillation is synchronized with the phase of lower-frequency oscillation. In other words, the strength of the higher-frequency oscillation depends on the timing of the lower-frequency oscillation. In this study, we considered high-frequency oscillation (2 to 500 Hz with a frequency resolution of 10 Hz) from LFP data corresponding to each of the spike electrodes and low-frequency oscillation (2 to 70 Hz with a frequency resolution of 4 Hz) from LFP data corresponding to each of the LFP electrodes.

To avoid edge artifacts, the entire data of each trial for every condition was first filtered across several passbands using a two-way, zero phase-lag, finite impulse response filter. The filter order was 3*r*, where r is the ratio of the sampling rate to the filter’s low-frequency cutoff. Subsequently, Hilbert transform was performed for each trial, to obtain phase from lower oscillation signal (2 to 70 Hz with a frequency resolution of 4 Hz) and amplitude from higher oscillation signal (2 to 500 Hz with a frequency resolution of 10 Hz). For every frequency bin of each condition, the phase or amplitude corresponding to the stimulus period (250 ms to 750 ms) of each trial was concatenated to make a single phase or amplitude signal. The coupling between the phase and amplitude signals in different frequency bins was calculated using the Kullback–Leibler Modulation Index (KLMI) (20), as described below.

For each phase and amplitude pair, the average amplitude of the amplitude-providing frequency in each phase bin of the phase providing frequency was computed. Here, the overall phases were divided into 18 bins (20° each). For each phase bin the average amplitude (p) was calculated as follows,

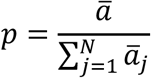

Where, 𝑎̅ is an average amplitude in each bin and 𝑁 is the number of bins (here, 18). The Shannon-entropy (*H*(*p*)) for each bin was calculated to compute the variation of amplitude in each phase bin.

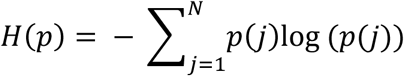

Then, the KLMI was computed by checking whether the distribution varied from uniform distribution using Kullback–Leibler distance.

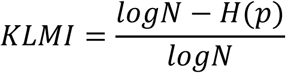

The peak angle was obtained by checking the maximum amplitude occurrence at each phase bin in average amplitude distribution (*p*).

To compute SFC, STA, and PAC, the evoked response was subtracted from LFP data and reconstructed LFP data using MP. Since we compared the effect of MP, which centers the data at zero, we subtracted the mean of LFP data before computing SFC, STA, and PAC, in addition to subtracting the evoked response. For visualization purposes, the frequency axis in all the plots is in log scale.

### Eliminating the sharp transients

#### Matching Pursuit (MP)

In this study, we primarily employed matching pursuit (31)(https://github.com/supratimray/MP.git) to eliminate the sharp transients from LFP data. We used a dictionary-based method with 500 Gabors or atoms. That is, we decomposed the entire data into 500 atoms and then removed atoms that had center frequency as zero corresponding to Gaussians or delta functions. Then, we reconstructed the signal with the remaining atoms. The reconstructed signal was used for computing SFC, STA, and PAC.

#### Neighboring LFP electrodes within 400 μm

For each spike electrode, we identified neighboring LFP electrodes within 400 μm. This resulted in 125 (for 38 spike electrodes) and 36 (for 17 spike electrodes) LFP electrodes for Monkey 1 and Monkey 2, respectively. SFC, STA, and PAC were computed between each spike electrode and its neighboring LFP electrode.

## Declaration of interests

The authors declare no competing financial interests.

## Funding disclosure and Acknowledgements

This work was supported by Wellcome Trust/DBT India Alliance (Senior fellowship IA/S/18/2/504003) to SR and Pratiksha Postdoctoral Fellowship to PP.

## Data and Code availability

Codes used in analysing the data and generating the figures associated with the study are available on GitHub (https://github.com/poojaprabhu9/CrossFrequencyCouplingProject.git).

---

**Supplementary Figure 1:**
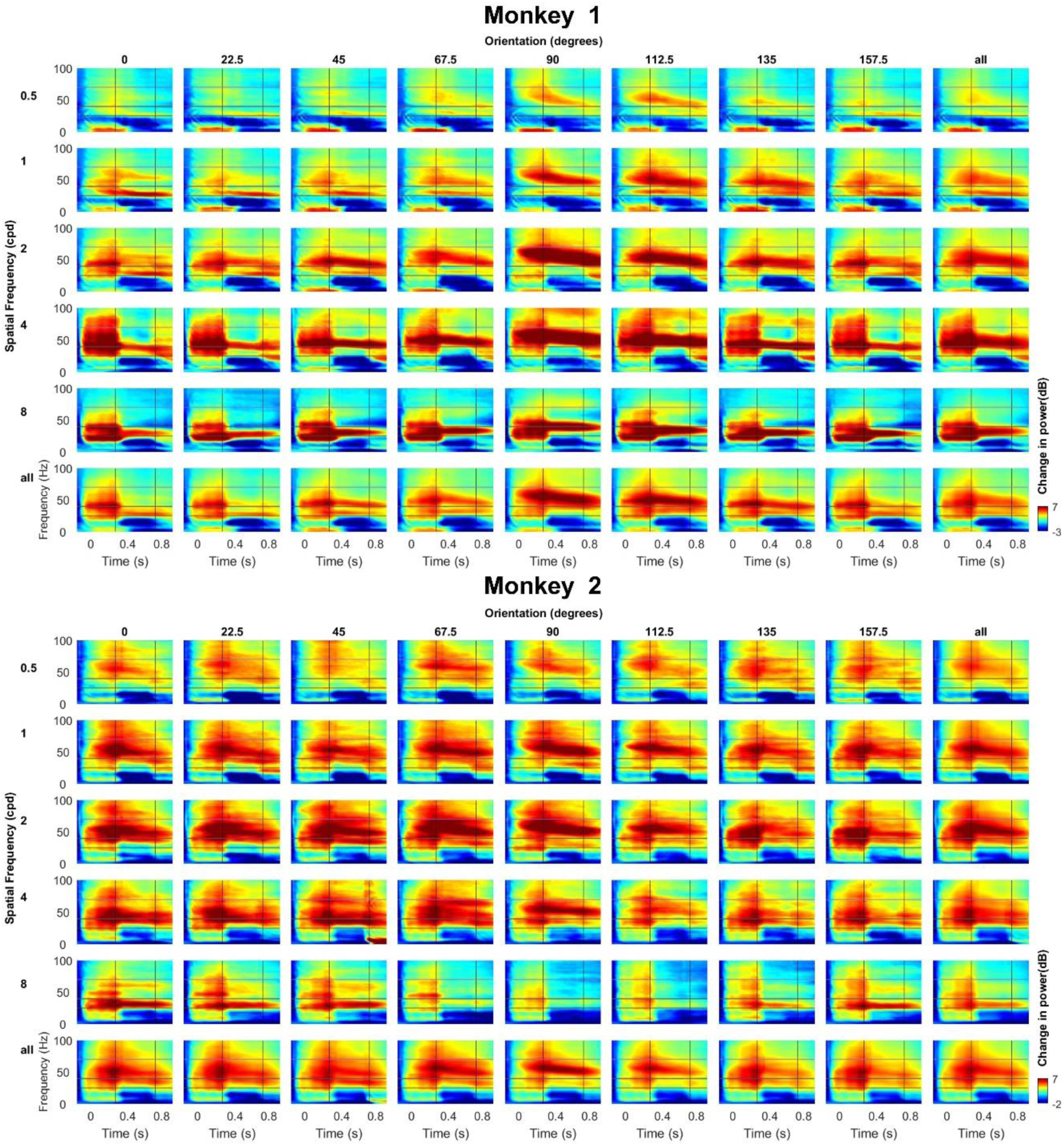
Normalized time-frequency plots across different spatial orientations (in rows) and orientations (in columns) for Monkey 1 and Monkey 2, averaged across LFP electrodes (N=38 for Monkey 1 and N=17 for Monkey 2). These show two gamma bands (slow gamma (25-40 Hz; brown horizontal line) and fast gamma (40-70 Hz; violet horizontal line), which were tuned to both orientation and spatial frequency. Stimulus was shown between 0 to 0.8 seconds, and oscillatory responses were most prominent after the initial transient period between 0.25 to 0.75 s (vertical lines), which was chosen as the “stimulus period” for analysis.

**Supplementary Figure 2:**
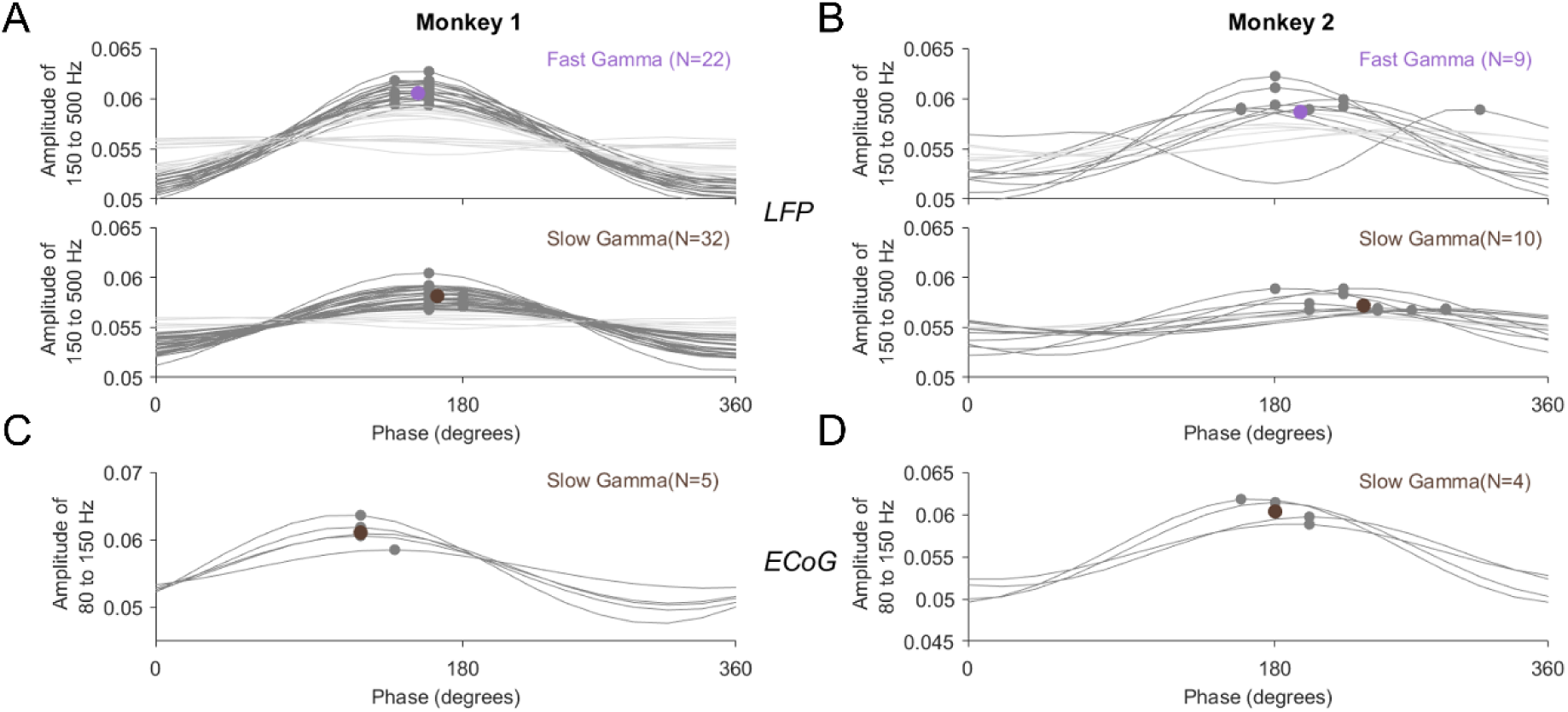
The phase versus mean amplitude distribution in LFP and ECoG. The trough of the low-frequency rhythm has a phase of 180 degrees (A-B) The distribution of the mean amplitude of the 150-500 Hz filtered LFP over the phases of the 40-70 Hz filtered LFP (top panel) and amplitude of 80-150 Hz filtered LFP over the phases of the 25-40 Hz filtered LFP (bottom panel). Each trace corresponds to one electrode. The trace is shown in dark gray if the PAC was significant using a bootstrap test or light gray otherwise. Mean of the phases of significant electrodes is shown as a dot. C-D) Same for ECoG data for the slow-gamma condition.

## Notes

### Competing Interest Statement

The authors have declared no competing interest.

